# Genomic Insights into the Population History and Adaptive Traits of Latin American Criollo Cattle

**DOI:** 10.1101/2023.09.10.556512

**Authors:** James A. Ward, Said I. Ng’ang’a, Imtiaz A. S. Randhawa, Gillian P. McHugo, John F. O’Grady, Julio M. Flórez, John A. Browne, Ana M. Pérez O’Brien, Antonio J. Landaeta-Hernández, José F. Garcia, Tad S. Sonstegard, Laurent A. F. Frantz, Michael Salter-Townshend, David E. MacHugh

## Abstract

Criollo cattle, descendants of animals brought by Iberian colonists to the Americas, have been subject to centuries of natural and human-mediated selection in novel tropical agroecological zones. Consequently, these breeds have evolved distinct characteristics such as resistance to disease and exceptional heat tolerance. In addition to European taurine (*Bos taurus*) ancestry, it has been proposed that gene flow from African taurine and Asian indicine (*Bos indicus*) cattle has shaped the ancestry of Criollo cattle. In this study, we analysed Criollo breeds from Colombia and Venezuela using whole-genome sequencing (WGS) and single-nucleotide polymorphism (SNP) array data to examine population structure and admixture at high resolution. Analysis of genetic structure and ancestry components provided evidence for African taurine and Asian indicine admixture in Criollo cattle. In addition, using WGS data, we detected selection signatures associated with a myriad of adaptive traits, revealing genes linked to thermotolerance, reproduction, fertility, immunity, and distinct coat and skin coloration traits. This study underscores the remarkable adaptability of Criollo cattle and highlights the genetic richness and potential of these breeds in the face of climate change, habitat flux, and disease challenges. Further research is warranted to leverage these findings for more effective and sustainable cattle breeding programmes.

## INTRODUCTION

The term “Criollo”, with origins in the Portuguese “Crioulo”, historically distinguished people born in the New World from those native to Iberia and was subsequently extended to livestock (cattle, sheep, horses, and goats). European livestock production in the Americas can be traced back to Columbus’ second voyage in 1493, where cattle, among other animals, sourced from La Gomera in the Canary Islands were brought to the island of Hispaniola [1–3]. Adoption and growth of cattle agriculture on Hispaniola was rapid; however, the spread of animals to the South American continent was more gradual. This was documented by early colonial accounts, illustrating a transformative ecological and economic process that started in the Caribbean, then Mexico, and eventually reaching the Ilanos of Colombia and Venezuela, where by 1600, cattle ranching had become a significant industry [4, 5]. Over the ensuing five centuries, increasing numbers of cattle were exported to North and South America particularly as the process of European colonization intensified during the 18^th^ and 19^th^ centuries [6]. Evidence for a similar Iberian origin narrative has been described for sheep [7], pigs [8], and horses. For example, recent work has shown that historical and extant North American horse populations exhibit pronounced genetic affinities with Iberian horse populations [9]. The ancestry of modern South American cattle has also been shaped by indicine (zebu) cattle (*Bos indicus*) that were directly introduced from South Asia—primarily present-day India and Pakistan [10].

The extent of contributions from cattle outside Europe and the Indian subcontinent to the South and Central American cattle gene pools remains uncertain. Modern Iberian cattle populations, whose ancestors likely contributed the bulk of Latin American Criollo cattle ancestry have an African taurine cattle genomic component [11–13]. In parallel to this, it has also been posited that African cattle may have been directly introduced into South America alongside imports from Europe [14]. This hypothesis is supported by the presence of the T1-c mtDNA haplotype at a frequency of 8% in Criollo cattle; T1-c is prevalent in African cattle and found at low frequencies in Iberian populations [15]. It is also supported by recent findings of colonial era mitogenomes which reveal the presence of African haplogroups in 17^th^ century cattle from Mexico [16]. Recent work suggests that the degree of African ancestry in Criollo cattle depends on the breed. Notably, the Guadeloupe breed, indigenous to the Caribbean, traces almost 35% of its genetic ancestry back to African taurine cattle [17].

The cattle that were brought to the Americas from Iberia in the 15^th^ and 16^th^ centuries were adapted to Mediterranean agroecologies and, over the following centuries, these populations have evolved adaptations to tropical and arid environments, emerging as distinctive breeds with unique heat tolerance and disease resistance traits [12, 18–21]. However, Criollo cattle are undervalued in modern production systems and face gradual replacement by more productive commercial breeds, which is eroding indigenous cattle genetic resources in Latin America [22]. Using genome-scale data, this study aims to evaluate *B. taurus* and *B. indicus* ancestry and detect historical gene flow from African cattle into the Blanco Orejinegro, Hartón del Valle, and Limonero Criollo breeds. Additionally, we use high-resolution analyses of genetic structure, and selection signatures to provide new information about the microevolutionary histories of these populations.

## MATERIALS AND METHODS

### Cattle samples, library preparation, high-throughput sequencing, and variant calling

In total, 34 Criollo cattle were used to generate high-quality de novo whole-genome sequence (WGS) data sets. Coat hair follicles obtained from 24 Colombian and Venezuelan cattle and semen straws from 10 Colombian AI bulls were used to obtain genomic DNA using standard commercial procedures for livestock DNA extraction and purification (Weatherbys Scientific, Co. Kildare, Ireland). The WGS data generation was performed by sequencing to a mean depth of 20× using two commercial service providers (Novogene, Cambridge, UK and Macrogen, Seoul, South Korea). Figure 1 displays the geographical origins of the three main Criollo breeds in this study.

**Figure 1.**
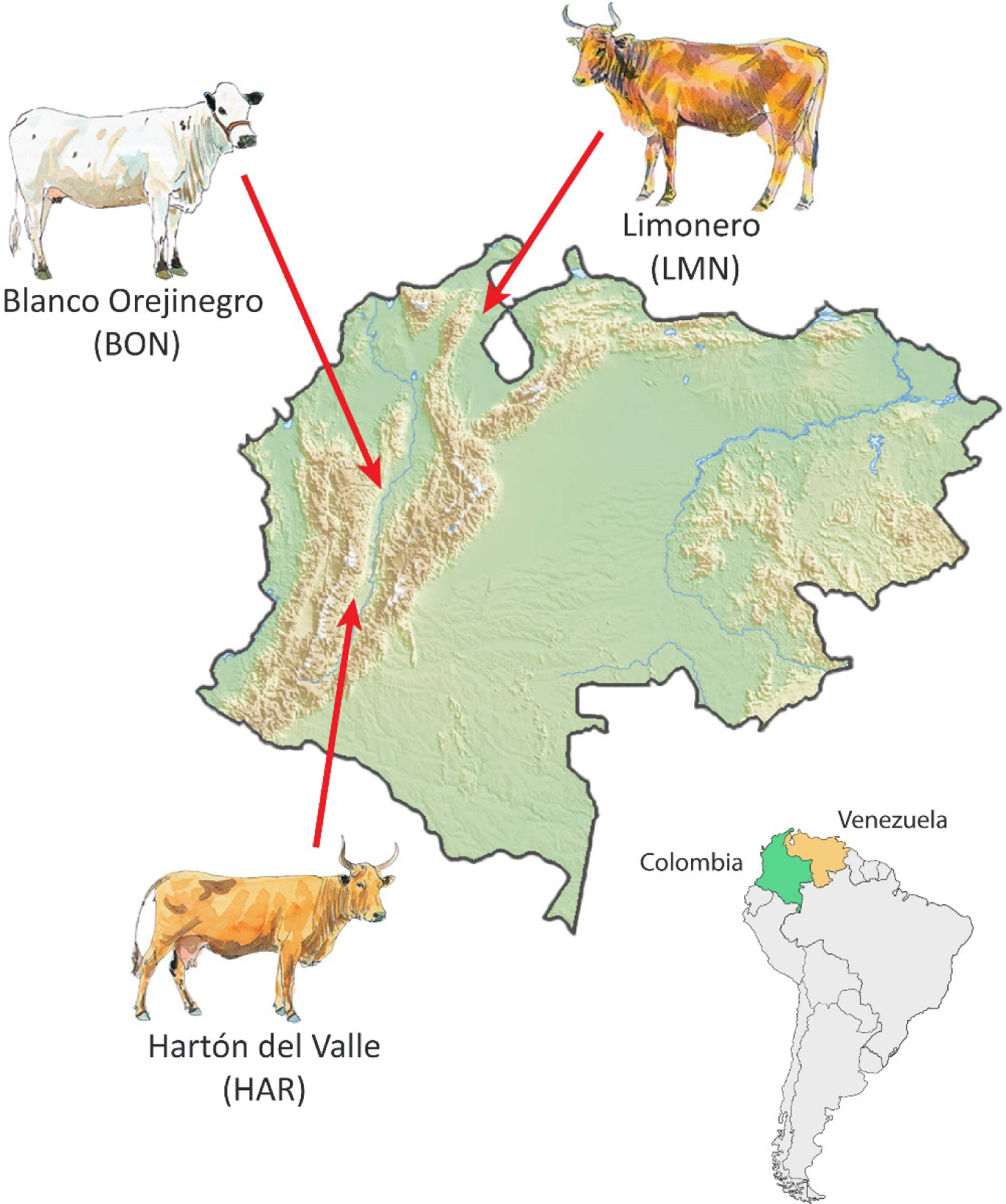
Approximate geographical origins of the animals from three Criollo breeds that were used to generate new whole-genome sequence data. Map modified from original by Mapswire (https://mapswire.com) (Creative Commons Attribution 4.0 International License CC-BY 4.0). Individual cattle art images modified from Felius [27] with permission of the author.

Publicly available WGS data was also assembled for 37 European taurine, 13 American indicine, and 13 American Criollo cattle. Sequence reads were mapped to the bovine reference genome (ARS-UCD1.2) [23] using the default parameters for bwa-mem2 v.2.2.1 [24]. Samtools v.1.13 [25] was used to sort bam files and create index files. Potential PCR duplicates were identified using the “MarkDuplicates” function of Picard v.2.27.1 (http://broadinstitute.github.io/picard). The “BaseRecalibrator” and “ApplyBQSR” functions of the Genome Analysis Toolkit (GATK) v.4.2.6.1 [26] were used to perform base quality score recalibration.

Candidate SNPs were called from the bam files using the “HaplotypeCaller” function in GATK with the “--emit-ref-confidence GVCF” option. Individual GVCF files were merged using the “GenomicsDBImport” function. SNPs from the genomics database were called and selected using “GenotypeGVCFs” and the “SelectVariants” function. The “VariantFiltration” function of GATK was used to avoid possible false-positive calls according to GATK best practices: SNP clusters with “--cluster-size 3” and “--cluster-window-size 10”, quality by depth “QD <2”, read position rank sum test “ReadPosRankSum < -8”, phred-scaled variant quality score “QUAL < 30”, Fisher strand “FS > 60”, mapping quality “MQ < 40”, mapping quality rank sum test “MQRankSum < -12.5” and strand odds ratio “SOR > 3”. Triallelic SNPs and those with a minor allele frequency < 0.01 and/or missing genotype rates over > 0.1 were filtered out. The SNPs remaining in each WGS data set were then annotated according to their positions using SnpEff [28]. The combined panel of 97 cattle samples with WGS data are detailed in electronic supplementary material, table S1.

### BovineHD 777k array data

For broader comparative population genomics analyses, publicly available BovineHD 777k SNP array data were used. These were 91 European taurine cattle representing two breeds, 82 Asian and American indicine cattle representing four breeds [29, 30], 45 Iberian cattle representing 13 breeds [31], and 31 African taurine cattle representing two breeds [29, 32]. In addition, 12 Criollo cattle representing the Senepol breed were obtained from the WIDDE database [33]. SNP positions were updated from the UMD3.1 assembly [34] to ARS-UCD1.2 following Riggio et al. [35]. Subsequent processing of these data was conducted using PLINK v1.90b6.25 [36] and R version 4.1.3 [37]. The combined panel of 261 cattle samples with BovineHD 777k SNP data are detailed in electronic supplementary material, table S2.

### Population differentiation, genetic structure, and demographic analyses

Principal component analysis (PCA) for all animals (WGS and BovineHD genotype data) were performed using PLINK v1.90b6.25 and the results were plotted using ggplot2 v3.4.0 [38] in the R v4.1.3 environment. The genetic structure of each population was visualized using ADMIXTURE v1.3.0 [39] with *K* = 3–6 unsupervised modelled ancestries and ggplot2 v3.4.0. Demographic modelling of current and historical effective population size (*N*_e_) trends was conducted using the SNeP software tool [40] with WGS data for the Blanco Orejinegro, Hartón del Valle, Limonero, N’Dama, and Holstein populations.

### Detection of historical admixture

Using ADMIXTOOLS [41] with default parameters, we calculated *f*_3_ statistics using the combined WGS and BovineHD genotype data set to test for evidence that three Criollo breeds (Blanco Orejinegro, Hartón del Valle, and Limonero) and three Iberian breeds (Limia, Pajuna, and Sayaguesa) are derived from the admixture of two sets of populations (European taurine populations and indicine cattle, and European taurine populations and African taurine cattle). A significant negative *f*_3_ value is considered evidence of historical admixture in the target population (Criollo or Iberian). For visualization of the *f*_3_ statistics, we plotted the *f*_3_ values against the population trios with error bars indicating the standard errors using ggplot2 v3.4.0 [38].

### Test statistics for detection of selection signatures

For the selection signatures analysis, only the WGS data set was used, which included approximately 9.7 million SNPs genotyped in 97 cattle. SNP-by-SNP *F*_ST_ values were calculated for each Criollo breed versus a panel of Holstein cattle using VCFtools v.0.1.16 [42]. XP-EHH statistics were calculated using the program selscan v.1.2.0 [43] with default parameters, where each Criollo breed was also compared to the Holstein panel. The unstandardized XP-EHH values were then normalized using the --norm function in selscan. We calculated the ΔSAF (directional change in selected allele frequency) using the --freq function of VCFtools to generate allele frequencies, then using a custom R script we calculated the ΔSAF for each Criollo breed versus the Holstein panel. Similarly to the XPEHH values, ΔSAF values were standardized to *Z* ∼ *N*(0,1) as described by Randhawa et al. [44].

### Implementing the composite selection signals (CSS) methodology

To obtain composite selection signals (CSS) statistics [44] for each Criollo breed, the *F*_ST_, XP-EHH, and ΔSAF statistics were combined by SNP ID (chromosome:position), which produced final sets of SNPs for the CSS analysis of 9,344,833, 9,465,007 and 9,660,647 million SNPs, for the Blanco Orejinegro, Hartón del Valle, and Limonero breeds, respectively. Following this, the MINOTAUR R package [45] was used to calculate CSS statistics, which were then averaged over 20 kb windows to reduce spurious signals. In addition, a signal was only considered significant if at least one SNP in the 0.1% of CSS scores was flanked by at least five other SNPs in the top 1% of CSS values [44, 46].

### Identification of genomic regions under selection

From the top 0.1% of SNPs identified, genomic intervals in the format chromosome:start-end were extracted and the BovineMine resource (v1.6) [47] was then used to find genes ± 500 kb from the start and end of these regions. The functional relevance of these CSS-derived gene sets was then evaluated using the scientific literature and the g:Profiler software tool [48], which was used to identify overrepresented gene ontology (GO) terms for each of the CSS-derived gene sets by breed. Following this, BovineMine v1.6 was used to systematically identify previously described quantitative trait loci (QTLs) for production, health, and welfare traits that are located within the CSS cluster regions for each breed.

## RESULTS

### Population differentiation, genetic structure, and admixture in Criollo cattle

The merged WGS and remapped BovineHD 777K BeadChip datasets yielded 302,012 shared SNPs for 308 individual animals. A PCA of these SNPs demonstrated that PC1 and PC2 accounted for 44.7% and 12.9% of the variation observed across the first 20 PCs, with PC1 representing the split between the *B. taurus* and *B. indicus* lineages and PC2 the split between the European and African taurine lineages (figure 2*a*). Genetic structure was analysed using ADMIXTURE, with cluster numbers (*K*) ranging from 3 to 6 (figure 2*b*). At *K* = 3, we recovered the three clusters observed in the PCA, which delineated European *B. taurus*, African *B. taurus*, and *B. indicus* ancestry components, respectively. The subsequent split at *K* = 4 captured an additional cluster within the European taurine populations that differentiated Jersey cattle. At *K* = 5 a Holstein cattle ancestry component was evident and at *K* = 6 a split was observed that divided *B. indicus* ancestry into two clusters.

**Figure 2.**
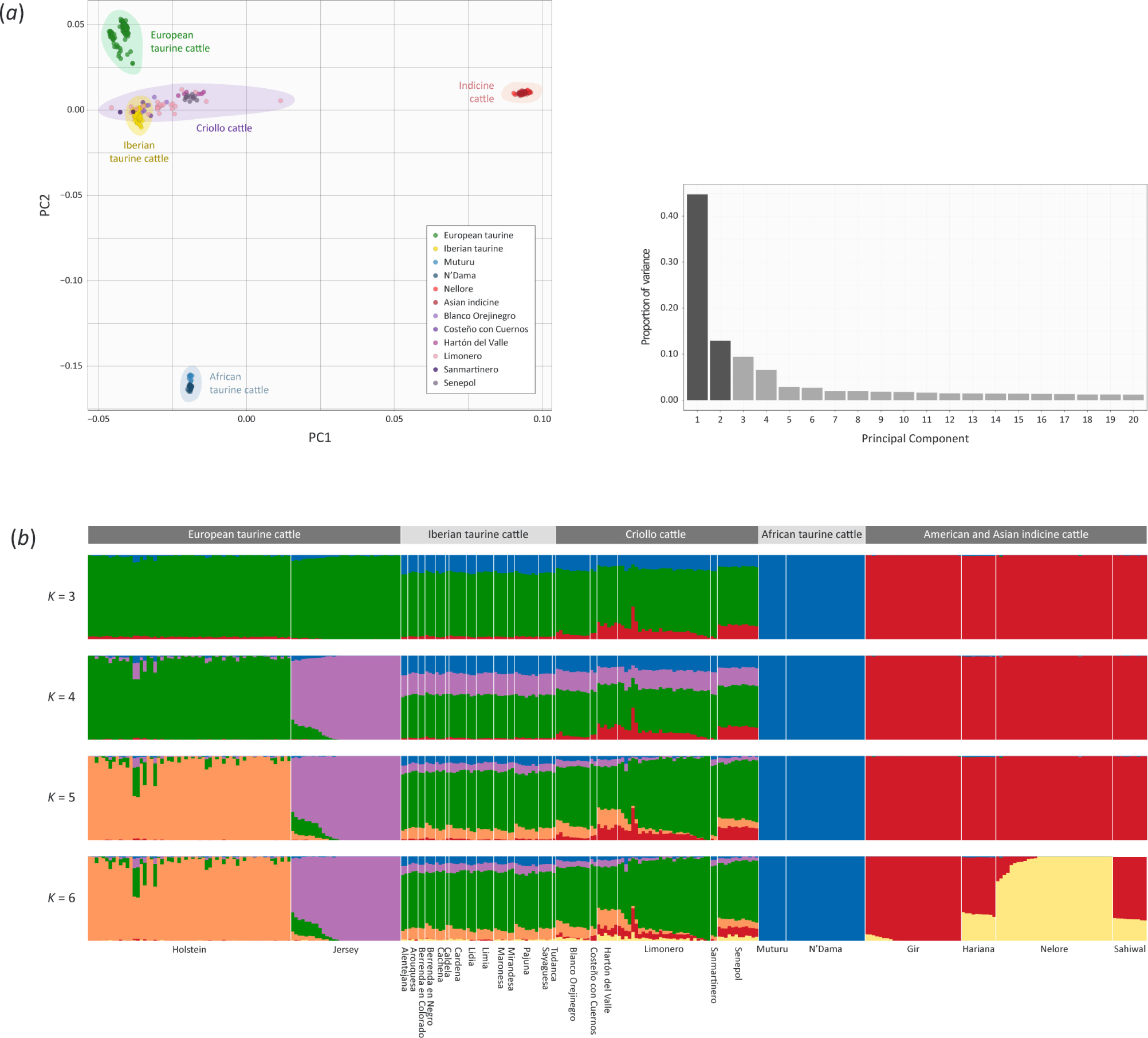
Genetic diversity and admixture in Criollo and comparator cattle populations. (*a*) Results of the principal component analysis (PCA) for 308 animals (47 from whole-genome sequence data and 261 from high-density SNP array data) with a shared set of 302,012 SNPs. The PCA plot shows the coordinates for each animal based on the first two principal components with the major breed groupings highlighted. Principal component 1 (PC1) differentiates the *Bos taurus* and *Bos indicus* lineages, whereas PC2 separates the African and European taurine groups. A histogram plot also shows the variance contributions for the first 20 PCs, with PC1 and PC2 accounting for 44.7% and 12.9% of this variance, respectively. (*b*) Unsupervised genetic structure plot for European and Iberian taurine, Criollo cattle, African taurine, and American and Asian indicine populations. Results for ancestry clusters ranging from *K* = 3–6 are shown.

Further examination of the PCA and structure plots in figure 2*a* and 2*b* showed that the Criollo cattle samples cluster with the Iberian taurine cattle samples, all of which show relatively uniform levels of African taurine admixture at *K* = 3 and *K* = 4. In addition, the Senepol and Hartón del Valle Criollo breeds display a consistent and relatively uniform ancestry component from the *B. indicus* lineage, while the Blanco Orejinegro, Costeño con Cuernos, Limonero, and Sanmartinero breeds exhibit more heterogeneous patterns of indicine admixture. Conversely, *f*_3_ statistics [41] indicate that there is no evidence of admixture from either African taurine or indicine cattle in the Blanco Orejinegro and Limonero breeds when using any of the European taurine reference populations and African taurine or indicine breeds as the second reference (electronic supplementary material, figure S1*a* and *c*). However, there is some evidence of indicine admixture in the Hartón del Valle as the *f*_3_ statistics for all population trios that used any European taurine reference population with an indicine breed as the second reference population produced negative *f*_3_ values for this breed (electronic supplementary material, figure S1*b*). This is also evident from the PCA (figure 2*a*), and the structure plot (figure 2*b*).

### Selection signals across Criollo cattle breed genomes

The CSS test identified selection signals for the three Criollo cattle breeds that had WGS data and sufficient sample sizes (Blanco Orejinegro, Hartón del Valle, and Limonero). For each breed, the test highlighted several selection signals spanning multiple chromosomes. For the Blanco Orejinegro breed, the CSS test detected 31 cluster regions spanning 17 chromosomes, with 442 genes located within these regions and extended ±500 kb (figure 3*a*). Similarly, for the Hartón del Valle breed, the CSS test revealed 31 distinct clusters distributed over 17 chromosomes, encompassing 410 genes (figure 3*b*). Lastly, for the Limonero breed, the CSS test detected 38 cluster regions, also extending across 17 chromosomes, and that contained 451 genes in total (figure 3*c*). When these results were consolidated, 76 genes within selection signature regions are shared across the three breeds, and BTA18 was the chromosome with the largest number of these genes (28 genes; 36.8% of the total). The CSS cluster regions and the genes within these regions are detailed by breed in electronic supplementary material, tables S3 and S4).

**Figure 3.**
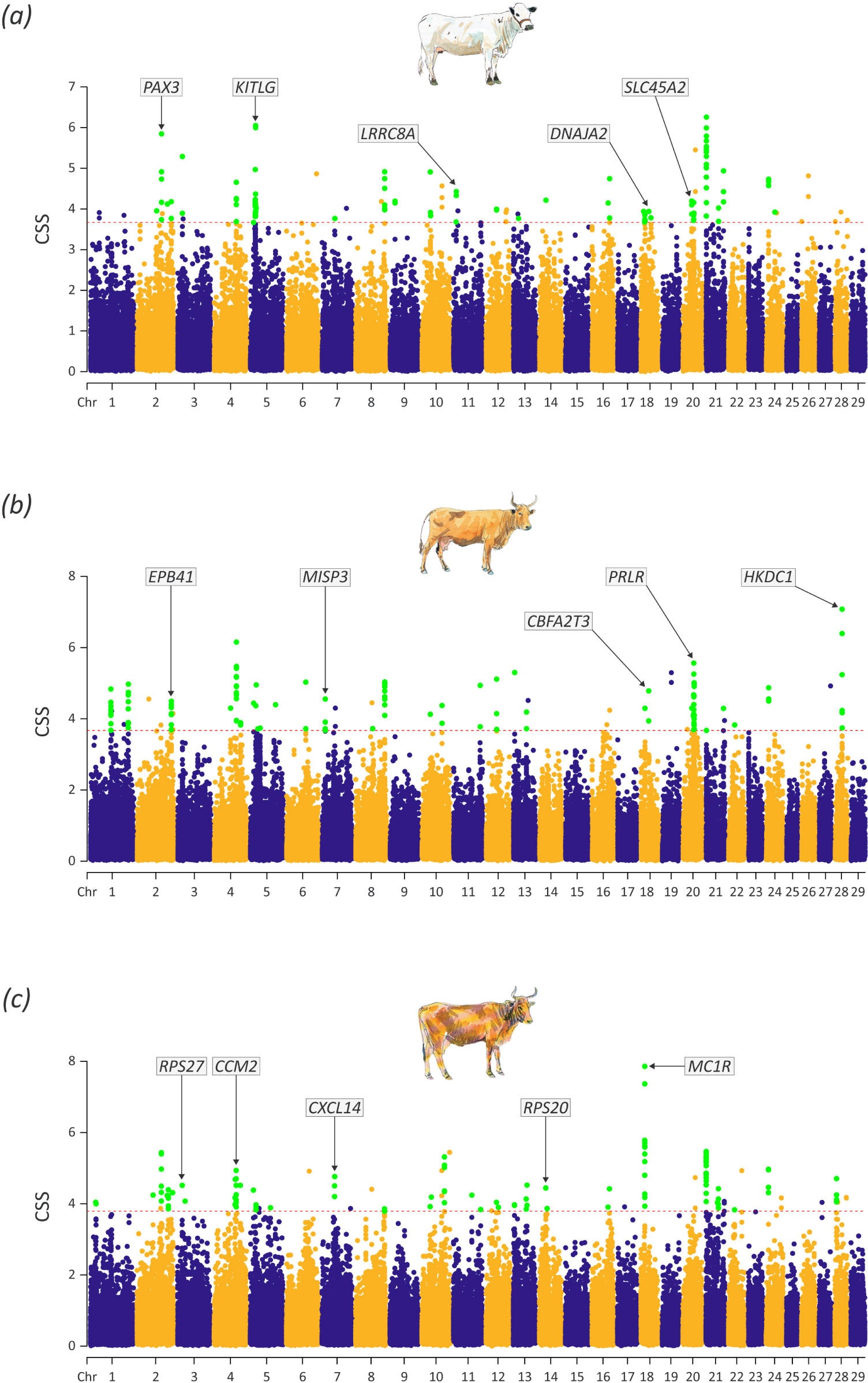
Chromosome-wide Manhattan plots of the smoothed composite selection signals (CSS) scores for Blanco Orejinegrio (*a*), Hartón del Valle (*b*) and Limonero (*c*). The dashed red lines indicate the genome-wide 0.1% thresholds of the empirical CSS scores. Green data points represent SNPs classified as significant and flanked by at least five additional SNPs among the top 1% of CSS scores. Notable candidate genes are indicated and discussed further in the text. Individual cattle art images modified from Felius [27] with permission of the author.

### Gene ontology (GO) term enrichment analysis

The functional significance of the genes within the CSS cluster regions for each breed was evaluated using GO term overrepresentation analysis with the g:Profiler software tool. After adjusting for multiple tests and using a *P*-value adjusted for the false discovery rate (FDR-*P*_adj._ < 0.05), overrepresented GO terms were catalogued for each breed. The results of the GO term overrepresentation analysis are shown in the electronic supplementary material, tables S5–S7 and figure 4. For both the Blanco Orejinegro (figure 4*a*) and Limonero (figure 4*b*) breeds, the GO terms related primarily to immunobiology, encompassing functions such as immunoglobulin receptor binding, phagocytosis, complement activation, and antigen binding. Conversely, for the Hartón del Valle breed (figure 4*c*), the predominant GO terms were associated with skin epidermis and keratinization processes.

**Figure 4.**
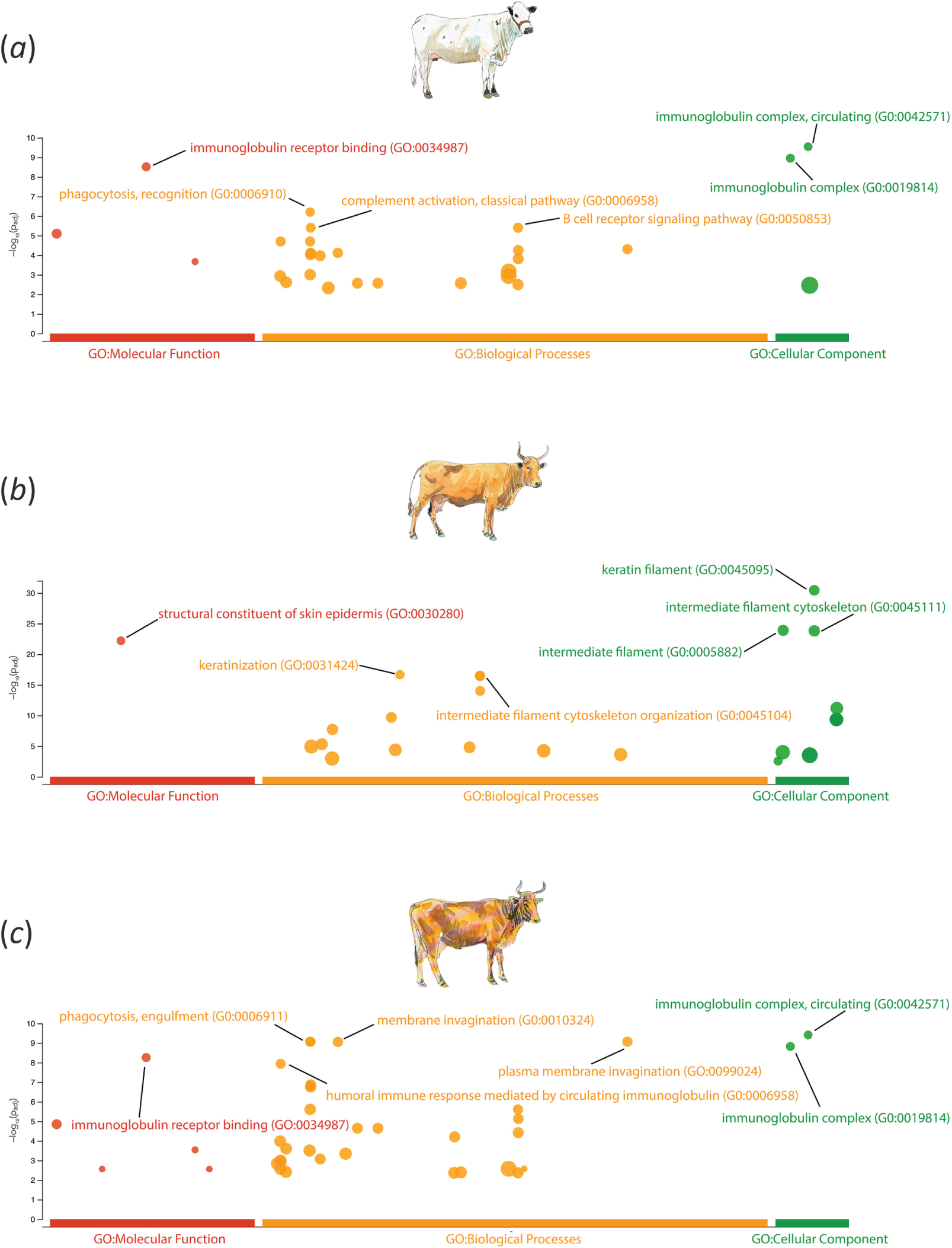
Gene ontology (GO) overrepresentation analysis results for the Blanco Orejinegrio (*a*), Hartón del Valle (*b*) and Limonero (*c*). The y-axis represents the negative log_10_ FDR-*P*_adj._ from the overrepresentation analysis. Individual cattle art images modified from Felius [27] with permission of the author.

### Identification of quantitative trait loci (QTLs) within CSS cluster regions

QTLs within the CSS cluster regions were systematically identified using BovineMine v1.6. The number of QTLs identified varied across the breeds: 519 in Blanco Orejinegro, 332 in Hartón del Valle, and 370 in Limonero. Interestingly, only 11 QTLs were common to all three breeds and a QTL associated with tick resistance [49] was particularly noteworthy. In the case of Blanco Orejinegro, most of the identified QTLs (291 out of 519) in the CSS cluster regions were associated with calving ease. Additionally, 46 QTLs were linked to milk protein percentage, 40 to milk yield, and 24 to coat texture. For the Limonero breed, there were 292 QTLs located in the CSS cluster regions related to calving ease; however, no other traits had more than five SNP markers among the QTLs. In the case of the Hartón del Valle breed, the most notable traits represented among the CSS cluster region QTLs were milk related. Specifically, 86 QTLs were associated with milk protein percentage, 45 with milk yield, and 35 with milk fat percentage. Additionally, 27 QTLs in Hartón del Valle were associated with coat texture. The results of the QTL search using BovineMine v1.6 are detailed in electronic supplementary material, table S8.

## DISCUSSION

### Population structure and admixture

The differentiation of cattle samples across PC1 of the PCA plot (figure 2*a*) reflects the evolutionary divergence between the *B. taurus* and *B. indicus* lineages, which occurred 150–500 kya [50, 51] The second division evident for PC2 in the PCA plot (figure 2*a*) differentiates the European and African taurine groups, a separation that has been well documented in several previous studies using genome-wide SNPs and other autosomal genetic markers [52–55]. These PCs are mirrored by the clusters observed at *K* = 3 in the structure analysis (figure 2*b*), which differentiate the European taurine, African taurine, and indicine groups and reveal patterns of shared ancestry among the major groups. Extending these observations further, complex patterns of admixture due to human-mediated and unmanaged crossbreeding between taurine and indicine cattle have also been described in cattle populations across several geographical regions, particular in sub-Saharan Africa [53–55].

Previous studies have noted African taurine autosomal admixture in Criollo cattle breeds [12, 17]. Results from the present study, which are captured in the PCA plot (figure 2*a*) and structure analysis (figure 2*b*) suggest this is also the case for the Criollo breed population samples analysed here (Blanco Orejinegro, Costeño con Cuernos, Hartón del Valle, Limonero, Sanmartinero, and Senepol), albeit at lower levels compared to that observed for Guadeloupe cattle [17]. Conversely, the positive *f*_3_ statistics observed for the three Criollo breeds examined using this method (Blanco Orejinegro, Hartón del Valle, and Limonero) do not support this (electronic supplementary material, figure S1*a*–*c*). It should be noted, however, that a positive *f*_3_ value does not necessarily imply a lack of admixture, since excessive genetic drift in the target population can obscure an admixture signal [41]. In this regard, demographic modelling of historical effective population sizes (*N*_e_ values) and comparison with the Holstein, a commercial dairy population with a markedly low *N*_e_ [56], indicates that founder effects and subsequent genetic drift may have impacted the Blanco Orejinegro, Hartón del Valle, and Limonero breeds (electronic supplementary material, figure S2).

For *B. indicus* ancestry, all the Latin American Criollo cattle populations examined here exhibit varying degrees of indicine admixture (figure 2*b*). Interestingly, however, the Blanco Orejinegro and Hartón del Valle both originate in Colombia but do not cluster together, which is presumably a consequence of the relatively modest indicine ancestry in the Blanco Orejinegro. The Limonero breed samples in this study largely cluster together except for four animals that exhibit a significant proportion of indicine ancestry. These results, taken together with the *f*_3_ statistics for the Blanco Orejinegro, Hartón del Valle, and Limonero (electronic supplementary material, figure S1*a–c*), and the histories of these breeds (the Blanco Orejinegro breed has been in a conservation programme since 1940 [57]), suggest that indicine gene flow has only had a long-term impact on breeds such as the Hartón del Valle which exhibits relatively uniform levels of indicine admixture compared to the Limonero, where it is more heterogeneous (figure 2*b*). This has implications for understanding the selection signatures that are described in the following sections: in some cases, the selection signals detected may be caused by adaptive introgression of *B. indicus* haplotypes.

Despite the inclusion of a diverse panel of European taurine, African taurine, Asian indicine, and admixed cattle in the design and validation of the BovineHD SNP array [58], we acknowledge that ascertainment bias could have influenced results from some of the analyses of population differentiation and genetic ancestry. However, we think that this is unlikely because ascertainment bias typically affects analyses such as selection signal detection that use individual SNP locus frequency-based statistics (e.g., *F*_ST_) substantially more than genome-wide multi-locus dimension reduction tools like ADMIXTURE and PCA [59–62]. Importantly, in this regard, we used only WGS data for the CSS analyses (figure 3 and electronic supplementary material, table S3).

### Genomic signatures of human-driven selection in Criollo cattle

Blanco Orejinegro are distinctive among Criollo cattle for their white coats, often speckled with black spots, and their skin is highly pigmented, as exemplified by their black ears. This coat-skin pattern also occurs in British White and White Park cattle [63], and for White Park, it has been demonstrated that this is caused by a duplication in the *KIT* gene and an aberrantly inserted *KIT* gene on BTA29 [64]. Additional genes that harbour cattle coat colour polymorphisms include *MC1R*, *TWIST2*, *MITF*, *PAX3*, *SLC45A2*, *COPA*, *TYR*, *TYRP1*, *KITLG*, and *ASIP*. For the Blanco Orejinegro breed, the genes *PAX3*, *SLC45A2*, and *KITLG* are in genome regions with evidence for directional selection based on the CSS results. Mutations in *PAX3* have been shown to result in a spectrum of phenotypic outcomes in horses from white spotting to a uniform white coat trait [65]. Similarly, mutations in the bovine *SLC45A2* gene have been shown to cause oculocutaneous albinism [66], while the *KITLG* gene has been associated with the roan coat type in cattle [67].

Interestingly, for the Hartón del Valle and Limonero breeds, *MC1R* was part of a BTA18 CSS cluster region. The protein receptor encoded by *MC1R* regulates tyrosinase levels in melanocytes, an enzyme critical to melanin synthesis. High concentrations of tyrosinase facilitate the production of eumelanin, contributing to darker brown or black shades, whereas lower concentrations drive the synthesis of phaeomelanin, yielding lighter red or yellow tones [68]. It is therefore noteworthy that Limonero and Hartón del Valle cattle generally possess a “bayo” or bay coat, which is a yellow to light brown/red colour [69, 70]. Though not in a CSS cluster region in the Blanco Orejinegro, a notable peak close to *MC1R* is present on BTA18. This gene has been observed to be under positive selection in *B. indicus* cattle resulting in a light grey to white coat in the Brahman and Nelore breeds [71]. The adaptive significance of functional polymorphisms at *MC1R* is that lighter coat colours reflect a significant portion of incident solar radiation, which makes these cattle more suited to tropical environments [72].

### Selection for heat tolerance in Criollo cattle

Criollo cattle breeds such as the Blanco Orejinegro, Hartón del Valle, and Limonero have evolved over five centuries to thrive in challenging tropical agroecologies. In the context of this environmental adaptation, a key finding for all three breeds is the presence of a CSS peak on BTA20 that encompasses the *PRLR* gene, most notably in the Blanco Orejinegro and Hartón del Valle breeds, where it was statistically significant. Mutations in this gene have been shown to cause the “slick” phenotype in in Criollo breeds such as the Blanco Orejinegro, Hartón del Valle, Limonero, Romosinuano, Costeño con Cuernos, Mexican Criollo Lechero Tropical, and also Criollo-influenced composite breeds such as the Carora and Senepol [73]. Of particular interest for understanding how convergent evolution can act in livestock populations is the existence of several different *PRLR* mutations that can produce the “slick” phenotype in Criollo cattle [73]. Mutations in the bovine *PRLR* gene can have major effects on the length and the structure of hair coats providing improved thermotolerance and concomitant increase on fertility and milk yields in cattle populations that inhabit dry and tropical conditions [18, 69, 74]. In addition, it has been shown that these mutations can act pleiotropically and cause other physiological changes [75].

Variation at other genes may also confer enhanced thermotolerance in Criollo cattle; for example, *MVD*, previously implicated as being under selection in North African cattle [76], was found in a BTA18 CSS cluster region for all three breeds. In addition, *CCM2* which regulates heart and blood vessel formation and integrity, is located in a Limonero BTA4 CSS cluster region and was also highlighted by Ben-Jemaa and colleagues in North African cattle [76]. This may represent an adaptation that increases heat transfer from the interior to the skin, which is also supported by the enhanced vascularisation of the dermis that has been documented in Limonero cattle [69]. Another gene, *SESN2*, located in a BTA2 CSS cluster detected in the Hartón del Valle and the Blanco Orejinegro breeds may also be associated with adaptation to tropical environments. This gene was observed to be downregulated in a functional genomics study of heat-stress responses in dermal fibroblast cells from indicine and indicine × taurine crossbred cattle [77].

The BTA11 Limonero CSS cluster region contains *FBXO4* that encodes a protein, which regulates body temperature through interactions with the heath shock protein HSPB6 (previously HSP20) [72, 78]. We also observed that *DNAJA2* was in a Blanco Orejinegro BTA18 CSS cluster region. This gene encodes a member of the DNAJ/HSP40 family of proteins and shows increased expression in peripheral blood from Holstein calves exposed to heat stress [79]. Similarly, *HIGD1A* (BTA22 CSS cluster) and *CBFA2T3* (a BTA18 CSS cluster distinct from that containing *DNAJA2*), encode proteins associated with cellular responses to hypoxia [80, 81], which could be indicative of adaptation to oxidative stress caused by high temperatures [82].

### Genomic signatures of selection for fertility and reproductive traits in Criollo cattle

Compared to taurine breeds from temperate zones, Criollo cattle are renowned for high fertility and excellent reproductive performance [21, 83] and using the CSS method we detected several genes associated with these traits. For example, in the Blanco Orejinegro breed a BTA7 CSS cluster region contains the *CATSPER3* gene, which encodes a sperm-specific ion channel protein directly linked to sperm motility and male fertility in many species, including cattle [84]. Another CSS cluster region in the Blanco Orejinegro on BTA20 contained *SPEF2*, which is involved in the formation and functionality of sperm flagella [85]. Additionally, the *CREM* gene, implicated in spermatogenesis, was located within a Blanco Orejinegro CSS cluster region on BTA13 [86]. Other genes related to reproductive physiology were located within CSS cluster regions for the Hartón del Valle breed including *HKDC1* on BTA28 that exhibits high expression in the testes [87], and *MISP3* on BTA7, which is implicated in spermatogenesis [88]. For the Limonero breed, CSS cluster regions included one on BTA10 that contains *FSIP1* a gene crucial for normal spermiogenesis and flagella development [89]. In addition, the *PTGES* gene within a Limonero CSS cluster region on BTA11 encodes an enzyme involved in the synthesis of prostaglandin E2 (*PGE2*), which is a signalling molecule with a crucial role in reproductive processes [90]. In parallel to this, the QTL analysis (electronic supplementary material, table S8) for the Blanco Orejinegro breed identified QTLs predominantly associated with reproduction, notably calving ease, and lactation. These findings corroborate previous results obtained using SNP array data for Blanco Orejinegro cattle [20, 91]. The results of the QTL analysis in the CSS cluster regions for the Limonero breed were similar to those for the Blanco Orejinegro breed.

### Genomic adaptations to infectious diseases in Criollo cattle

Comparable to reproductive and fertility traits, desirable disease resistance and tolerance traits have been highlighted as important features of Criollo cattle populations. However, empirical evidence for the genetic basis of these traits is often lacking; instead, much of the evidence base is anecdotal and derived from practical knowledge and experiences of veterinarians, farmers and breeders [70]. In the present study, however, we have begun to address this knowledge gap through identification of several immune genes in CSS cluster regions for the Blanco Orejinegro, Hartón del Valle, and Limonero breeds. For example, the *CBFA2T3* gene located in the BTA18 CSS cluster region detected for all three breeds and discussed in the context of hypoxia, has a role in the bovine immune response to mammary gland inflammation [92]. Similarly, *CXCL14*, located in a BTA7 CSS cluster region for the Blanco Orejinegro and Limonero breeds has also been implicated in the immune response to bovine mastitis caused by *Staphylococcus aureus* [93]. In addition, for the Limonero, a BTA11 CSS cluster region contains the *LRRC8A* gene, which is essential for development and function of T lymphocytes [94].

In tropical and subtropical regions, there is a significant prevalence of vector-borne haemoparasitic infections [95] that induce anaemia, fever, wasting, and reproductive dysfunction [96]. These diseases are primarily caused by eukaryotic parasites within the *Babesia* [97] and *Trypanosoma* [98] genera, and bacterial pathogens from the *Anaplasma* genus [99]. It is noteworthy, therefore, that the *EPB41* gene located within a Blanco Orejinegro and Hartón del Valle BTA2 CSS cluster region is associated with resistance to anaemia and trypanotolerance in African cattle [100]. Also, for the Blanco Orejinegro and Limonero, *RPS27* and *RPS20* are located in CSS cluster regions on BTA3 and BTA14, respectively and mutations in these genes can cause Diamond-Blackfan anaemia [101, 102]. Furthermore, in the Blanco Orejinegro breed, the *ZFPM1* gene within a BTA18 CSS cluster region, has been shown to play a role in cardiac and hematopoietic development. In a murine *Trypanosoma congolense* infection model, mice exhibiting decreased *ZFPM1* expression recovered more effectively from anaemia [103].

Gene GO term analysis in the Criollo CSS cluster regions revealed a marked overrepresentation of immunobiological processes for the Blanco Orejinegro and Limonero breeds, many of which overlapped between the two breeds (figure 4*a* and 4c and electronic supplementary material, tables S5 and S7). The statistically significant overrepresented biological process GO terms in the Blanco Orejinegro and Limonero included several terms related to intracellular pathogens, including *phagocytosis* (GO:0006909), *phagocytosis, engulfment* (GO:0006911), and *phagocytosis, recognition* (GO:0006910). It is therefore noteworthy that Blanco Orejinegro cattle have a documented resistance to brucellosis [104], a disease characterized by abortion and retention of the placenta, which is principally caused by *Brucella abortus* an intracellular bacterium that subverts phagocytic pathways and processes to enter and manipulate host cells [105].

## CONCLUSIONS

This study provides valuable insights into the genomic basis of microevolutionary change in Criollo cattle as they have adapted to the tropical environments of Latin America. Evidence of strong selective pressure was apparent, particularly for the distinct coat and skin coloration traits observed in these breeds, which are advantageous in cattle populations exposed to significant levels of incident solar radiation. Notably, we also discovered genomic selection signatures that may be associated with thermotolerance, again underscoring adaptation to hot climates. In addition, some of the selection signatures we identified align with documented fertility traits in Criollo cattle. Moreover, functional overrepresentation analysis revealed many genes related to immune function, which could reflect resilience to multiple infectious disease challenges. Taken together, our results, show the remarkable adaptability of Criollo cattle, which has been driven by natural and human-mediated selection and that underscores the genetic richness and value of these breeds for future breeding programmes.

## Acknowledgements

We thank Thomas Hall for advice on bioinformatics and computational genomics methodologies.

## Funding

J.A.W. was supported by Science Foundation Ireland (SFI) and Acceligen/Recombinetics Inc. through the SFI Centre for Research Training in Genomics Data Science under grant no. 18/CRT/6214. G.P.M. and D.E.M. were supported by an SFI Investigator Award (SFI/15/IA/3154). T.S.S. and D.E.M were supported by a United States Department of Agriculture (USDA) and Department of Agriculture, Food, and the Marine (DAFM) US-Ireland Research and Development Partnership grant (TARGET-TB, 17/RD/USROI/52).

## Notes

https://www.ebi.ac.uk/ena/browser/view/PRJEB65887

### Competing Interest Statement

The authors have declared no competing interest.

### Summary of Updates

We have redone some analyses with more appropriate comparator populations. We also rewritten the first part of the Introduction. In addition, some other small typographical errors/mistakes have been corrected.

