## Supplementary Material for "Genomic Insights into the Population History and Adaptive Traits of Latin American Criollo Cattle"

### Genomic Insights into the Population History and Adaptive Traits of Latin American Criollo Cattle

James A. Ward[1], Said I. Ng'ang'a[2,3], Imtiaz A. S. Randhawa[4], Gillian P. McHugo[1], John F. O'Grady[1], Julio M. Flórez[5,6], John A. Browne[1], Ana M. Pérez O'Brien[5], Antonio J. Landaeta-Hernández[7], José F. Garcia[6], Tad S. Sonstegard[5], Laurent A. F. Frantz[2,3], Michael Salter-Townshend[8], David E. MacHugh[1,9]

[1] Animal Genomics Laboratory, UCD School of Agriculture and Food Science, University College Dublin, Belfield, Dublin, D04 V1W8, Ireland.

[2] Palaeogenomics Group, Department of Veterinary Sciences, Ludwig Maximilian University, Munich D-80539, Germany.

[3] School of Biological and Chemical Sciences, Queen Mary University of London, London E1 4NS, United Kingdom.

[4] Faculty of Science, The University of Queensland, Gatton, QLD 4343, Australia

[5] Acceligen, 3388 Mike Collins Drive, Eagan, MN, 55121, USA.

[6] Department of Preventive Veterinary Medicine and Animal Reproduction, School of Agricultural and Veterinarian Sciences, São Paulo State University (Unesp), Jaboticabal, Brazil.

[7] Unidad de Investigaciones Zootécnicas, Facultad de Ciencias Veterinarias, Universidad del Zulia, Maracaibo, Venezuela.

[8] UCD School of Mathematics and Statistics, University College Dublin, Dublin, D04 V1W8, Ireland.

[9] UCD Conway Institute of Biomolecular and Biomedical Research, University College Dublin, Dublin, D04 V1W8, Ireland.

### Supplementary Figures

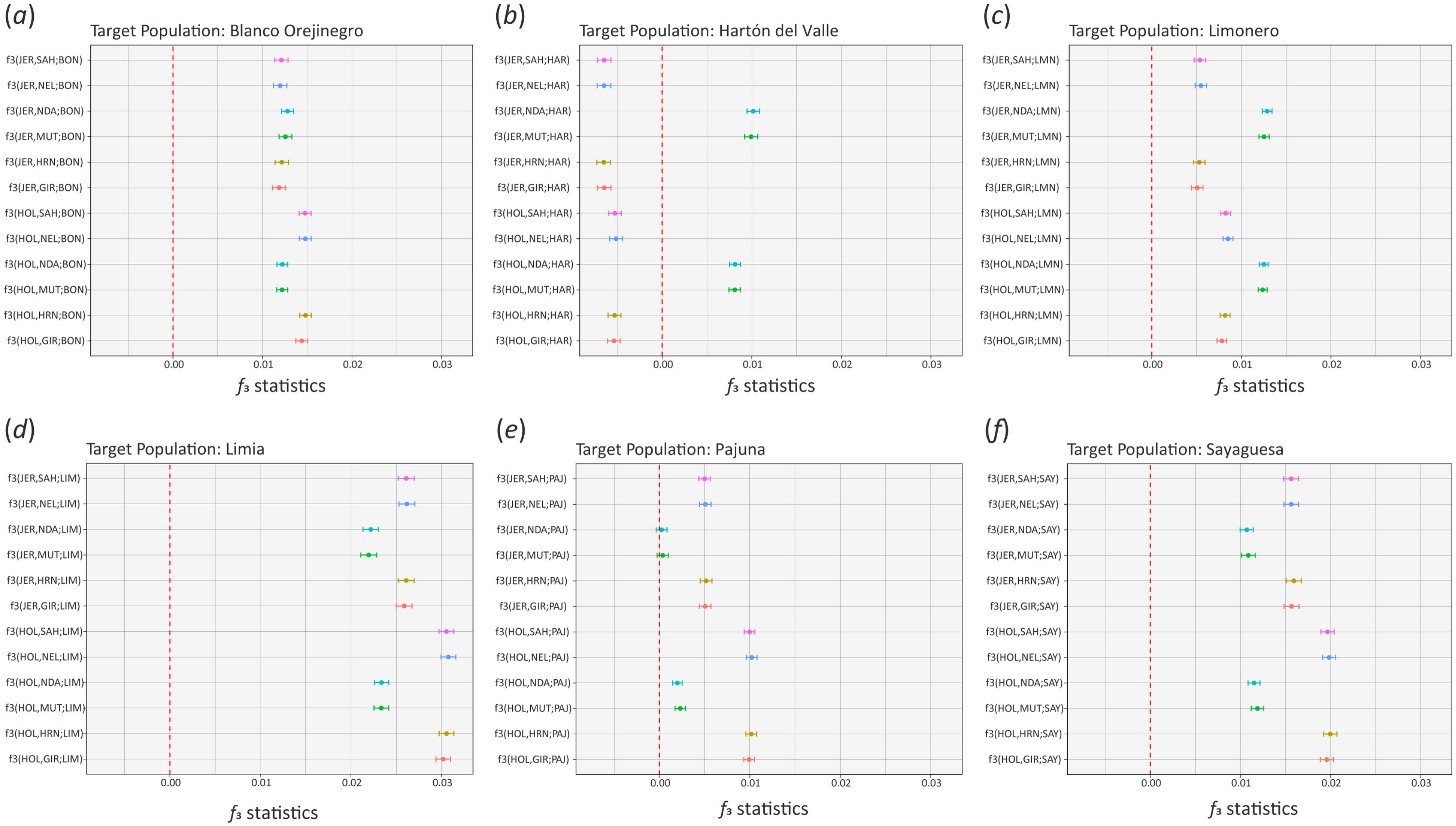


**Figure S1.** Three-population tests using the combined whole-genome sequence (WGS) and remapped BovineHD 777K BeadChip data for the Blanco Orejinegrio – BON (*a*), Hartón del Valle – HAR (*b*), and Limonero – LMN (*c*) Limia – LIM (*d*), Pajuna – PAJ (*e*), and Sayaguesa – SAY (*f*) as target populations. The reference populations used were two European taurine populations (Jersey – JER; and Holstein – HOL), four indicine breeds (Gir – GIR; Hariana – HRN; Nelore – NEL; and Sahiwal – SAH), and two African taurine populations (NDA – N'Dama; and Muturu – MUT). Each coloured data point shows the $f_3$ statistics value and the horizontal line represents plus or minus the standard error (±SE). Further details on the breeds/populations are also provided in Table S2.

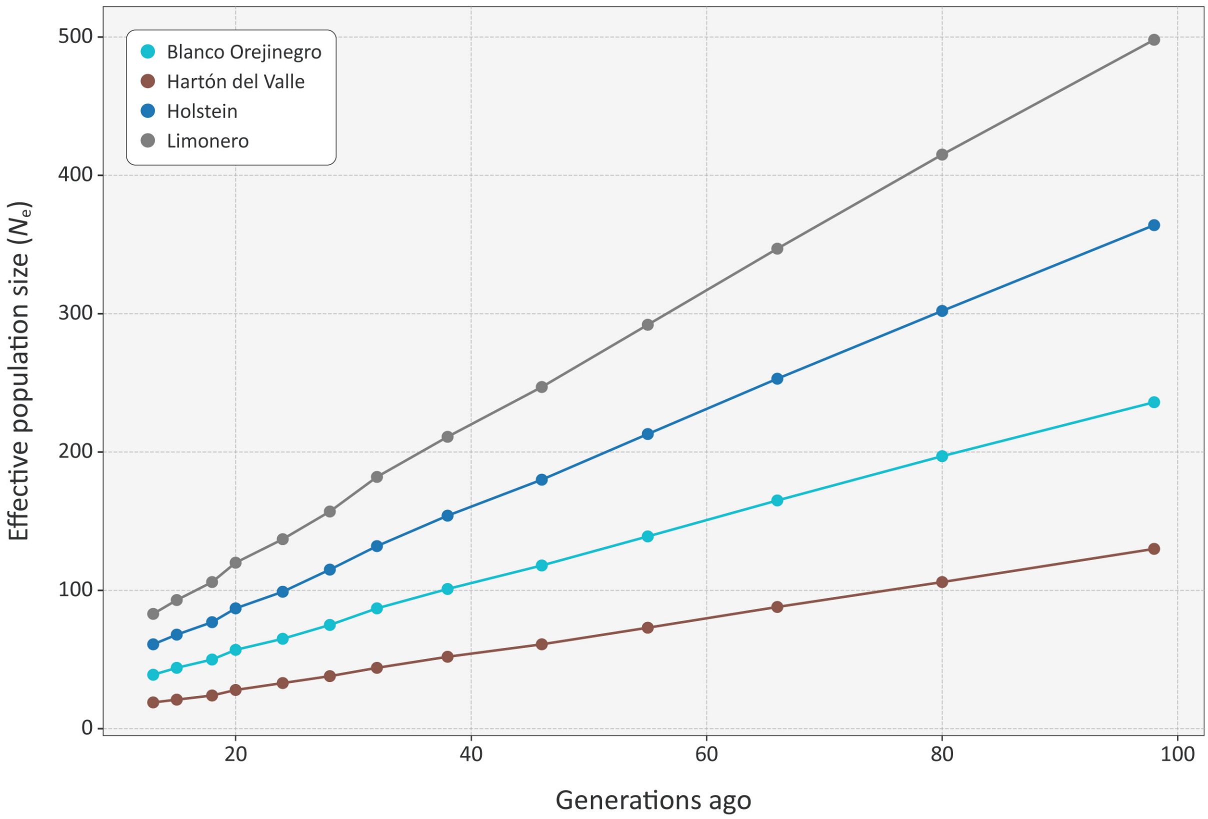


**Figure S2.** Modelled effective population size ($N_e$) values across time for the Blanco Orejinegro, Hartón del Valle, Limonero, and Holstein cattle breeds.

**Genomic Insights into the Population History and Adaptive Traits of Latin American Creole Cattle**

**Supplementary Tables**

**Table S1:** Cattle samples with whole-genome sequence (WGS) data.

| Sample sequence code | Breed/ pop. code | Lab code | Breed | Lineage | Phenotype | Source |
|---|---|---|---|---|---|---|
| SA01 | BON | BON01 | Blanco Orejinegro | American Creole | N/A | This study |
| SA02 | BON | BON02 | Blanco Orejinegro | American Creole | N/A | This study |
| SA03 | BON | BON03 | Blanco Orejinegro | American Creole | N/A | This study |
| SA04 | BON | BON04 | Blanco Orejinegro | American Creole | N/A | This study |
| SA05 | BON | BON05 | Blanco Orejinegro | American Creole | N/A | This study |
| SA06 | BON | BON06 | Blanco Orejinegro | American Creole | N/A | This study |
| SA07 | BON | BON07 | Blanco Orejinegro | American Creole | N/A | This study |
| SA08 | BON | BON08 | Blanco Orejinegro | American Creole | N/A | This study |
| SA09 | BON | BON09 | Blanco Orejinegro | American Creole | N/A | This study |
| SA10 | BON | BON10 | Blanco Orejinegro | American Creole | N/A | This study |
| SRR7170787 | CCC | CCC01 | Costeño con Cuernos | American Creole | N/A | PRJNA471656 |
| SRR7170788 | CCC | CCC02 | Costeño con Cuernos | American Creole | N/A | PRJNA471656 |
| G13 | HAR | HAR01 | Hartón del Valle | American Creole | N/A | This study |
| G16 | HAR | HAR02 | Hartón del Valle | American Creole | N/A | This study |
| G17 | HAR | HAR03 | Hartón del Valle | American Creole | N/A | This study |
| G20 | HAR | HAR04 | Hartón del Valle | American Creole | N/A | This study |
| G9 | HAR | HAR05 | Hartón del Valle | American Creole | N/A | This study |
| G18 | HAR | HAC01 | Hartón del Valle/Carora | American Creole | N/A | This study |
| G1 | LMN | LMN01 | Limonero | American Creole | N/A | This study |
| G10 | LMN | LMN02 | Limonero | American Creole | N/A | This study |
| G11 | LMN | LMN03 | Limonero | American Creole | N/A | This study |
| G12 | LMN | LMN04 | Limonero | American Creole | N/A | This study |
| G14 | LMN | LMN05 | Limonero | American Creole | N/A | This study |
| G19 | LMN | LMN06 | Limonero | American Creole | N/A | This study |
| G2 | LMN | LMN07 | Limonero | American Creole | N/A | This study |
| G21 | LMN | LMN08 | Limonero | American Creole | N/A | This study |

| Sample sequence code | Breed/ pop. code | Lab code | Breed | Lineage | Phenotype | Source |
|---|---|---|---|---|---|---|
| G22 | LMN | LMN09 | Limonero | American Creole | N/A | This study |
| G3 | LMN | LMN10 | Limonero | American Creole | N/A | This study |
| G5 | LMN | LMN11 | Limonero | American Creole | N/A | This study |
| G6 | LMN | LMN12 | Limonero | American Creole | N/A | This study |
| G7 | LMN | LMN13 | Limonero | American Creole | N/A | This study |
| G8 | LMN | LMN14 | Limonero | American Creole | N/A | This study |
| SRR6371016 | LMN | LMN15 | Limonero | American Creole | Slick coat | PRJNA422135 |
| SRR6371017 | LMN | LMN16 | Limonero | American Creole | Slick coat | PRJNA422135 |
| SRR6371018 | LMN | LMN17 | Limonero | American Creole | Slick coat | PRJNA422135 |
| SRR6371019 | LMN | LMN18 | Limonero | American Creole | Long coat | PRJNA422135 |
| SRR6371020 | LMN | LMN19 | Limonero | American Creole | Slick coat | PRJNA422135 |
| SRR6371021 | LMN | LMN20 | Limonero | American Creole | Slick coat | PRJNA422135 |
| SRR6371022 | LMN | LMN21 | Limonero | American Creole | Slick coat | PRJNA422135 |
| SRR6371023 | LMN | LMN22 | Limonero | American Creole | Slick coat | PRJNA422135 |
| SRR6371024 | LMN | LMN23 | Limonero | American Creole | Normal coat | PRJNA422135 |
| G23 | LAR | LAR01 | Limonero/Romosinuano | American Creole | N/A | This study |
| G24 | LAS | LAS01 | Limonero/Senepol | American Creole | N/A | This study |
| G4 | LAS | LAS02 | Limonero/Senepol | American Creole | N/A | This study |
| SRR7170789 | SAM | SAM01 | San Martinero | American Creole | N/A | PRJNA471656 |
| SRR7170790 | SAM | SAM02 | San Martinero | American Creole | N/A | PRJNA471656 |
| G25 | UKN | UKN01 | Unknown Creole breed | American Creole | N/A | This study |
| SAMN05788494 | AMI | BRA01 | Brahman | American Indicine | N/A | PRJNA343262 |
| SAMN05788495 | AMI | BRA02 | Brahman | American Indicine | N/A | PRJNA343262 |
| SAMN05788496 | AMI | BRA03 | Brahman | American Indicine | N/A | PRJNA343262 |
| SAMN05788512 | AMI | GIR01 | Gir | American Indicine | N/A | PRJNA343262 |
| SAMN05788513 | AMI | GIR02 | Gir | American Indicine | N/A | PRJNA343262 |
| SAMN05788514 | AMI | GIR03 | Gir | American Indicine | N/A | PRJNA343262 |
| SAMN05788515 | AMI | GIR04 | Gir | American Indicine | N/A | PRJNA343262 |

| Sample sequence code | Breed/ pop. code | Lab code | Breed | Lineage | Phenotype | Source |
|---|---|---|---|---|---|---|
| SAMN05788520 | AMI | NEL01 | Nelore | American Indicine | N/A | PRJNA343262 |
| SAMN05788521 | AMI | NEL02 | Nelore | American Indicine | N/A | PRJNA343262 |
| SAMN05788522 | AMI | NEL03 | Nelore | American Indicine | N/A | PRJNA343262 |
| SAMN05788523 | AMI | NEL04 | Nelore | American Indicine | N/A | PRJNA343262 |
| SAMN05788524 | AMI | NEL05 | Nelore | American Indicine | N/A | PRJNA343262 |
| SAMN05788525 | AMI | NEL06 | Nelore | American Indicine | N/A | PRJNA343262 |
| SRR4477870 | EUT | ANG01 | Angus | European Taurine | N/A | PRJNA343262 |
| SAMN05788492 | EUT | HOL01 | Holstein | European Taurine | N/A | PRJNA343262 |
| SAMN05788508 | EUT | HOL02 | Holstein | European Taurine | N/A | PRJNA343262 |
| SRR4279977 | EUT | HOL03 | Holstein | European Taurine | N/A | PRJNA343262 |
| SRR4279978 | EUT | HOL04 | Holstein | European Taurine | N/A | PRJNA343262 |
| SRR4279979 | EUT | HOL05 | Holstein | European Taurine | N/A | PRJNA343262 |
| SRR4280060 | EUT | HOL06 | Holstein | European Taurine | N/A | PRJNA343262 |
| SRR8587906 | EUT | HOL07 | Holstein | European Taurine | N/A | PRJNA343262 |
| SRR8587907 | EUT | HOL08 | Holstein | European Taurine | N/A | PRJNA343262 |
| SRR8587979 | EUT | HOL09 | Holstein | European Taurine | N/A | PRJNA343262 |
| SRR8587980 | EUT | HOL10 | Holstein | European Taurine | N/A | PRJNA343262 |
| SRR8587981 | EUT | HOL11 | Holstein | European Taurine | N/A | PRJNA343262 |
| SRR934405 | EUT | HOL12 | Holstein | European Taurine | N/A | PRJNA210521 |
| SRR934406 | EUT | HOL13 | Holstein | European Taurine | N/A | PRJNA210521 |
| SRR934407 | EUT | HOL14 | Holstein | European Taurine | N/A | PRJNA210521 |
| SRR934408 | EUT | HOL15 | Holstein | European Taurine | N/A | PRJNA210521 |
| SRR934409 | EUT | HOL16 | Holstein | European Taurine | N/A | PRJNA210521 |
| SAMN05788516 | EUT | JER01 | Jersey | European Taurine | N/A | PRJNA343262 |
| SAMN05788518 | EUT | JER02 | Jersey | European Taurine | N/A | PRJNA343262 |
| SRR3497161 | EUT | JER03 | Jersey | European Taurine | N/A | PRJNA318089 |
| SRR3497162 | EUT | JER04 | Jersey | European Taurine | N/A | PRJNA318089 |
| SRR3497451 | EUT | JER05 | Jersey | European Taurine | N/A | PRJNA318089 |

| Sample sequence code | Breed/ pop. code | Lab code | Breed | Lineage | Phenotype | Source |
|---|---|---|---|---|---|---|
| SRR3497462 | EUT | JER06 | Jersey | European Taurine | N/A | PRJNA318089 |
| SRR3497464 | EUT | JER07 | Jersey | European Taurine | N/A | PRJNA318089 |
| SRR8426539 | IBT | LIM01 | Limia | European Taurine | N/A | PRJNA514237 |
| SRR8426540 | IBT | MAR01 | Maronesa | European Taurine | N/A | PRJNA514237 |
| SAMN10721584 | EUT | PAJ01 | Pajuna | European Taurine | N/A | PRJNA514237 |
| SAMN10721579 | EUT | SAY01 | Sayaguesa | European Taurine | N/A | PRJNA514237 |
| SRR10752679 | EUT | SRH01 | Shorthorn | European Taurine | N/A | PRJNA343262 |
| SRR10752680 | EUT | SRH02 | Shorthorn | European Taurine | N/A | PRJNA343262 |
| SRR10752681 | EUT | SRH03 | Shorthorn | European Taurine | N/A | PRJNA343262 |
| SRR10752682 | EUT | SRH04 | Shorthorn | European Taurine | N/A | PRJNA343262 |
| SRR10752683 | EUT | SRH05 | Shorthorn | European Taurine | N/A | PRJNA343262 |
| SRR10752684 | EUT | SRH06 | Shorthorn | European Taurine | N/A | PRJNA343262 |
| SRR10752677 | EUT | SIM01 | Simmental | European Taurine | N/A | PRJNA343262 |
| SRR10752700 | EUT | SIM02 | Simmental | European Taurine | N/A | PRJNA343262 |
| SRR10752701 | EUT | SIM03 | Simmental | European Taurine | N/A | PRJNA343262 |

**Table S2:** Cattle samples with BovineHD 777k SNP data [Excel file – Ward_et_al.(2023)_Table_S2.xlsx]

**Table S3:** Composite selection signatures (CSS) cluster regions and genes for all breeds [Excel file – Ward_et_al.(2023)_Table_S3.xlsx]

**Table S4:** Composite selection signatures (CSS) genes and functional inference.

| Breed | Associated trait | Gene | Functional inference | References |
|---|---|---|---|---|
| Blanco Orejinegro | Coat colour | *PAX3* | Mutations in *PAX3* cause a spectrum of white spotting to entirely white. | [1] |
| Blanco Orejinegro | Coat colour | *SLC45A2* | Mutations in *SLC45A2* cause oculocutaneous albinism. | [2] |
| Blanco Orejinegro | Coat colour | *KITLG* | *KITLG* is linked to the roan coat type. | [3] |
| Limonero, Hartón del Valle | Coat colour | *MC1R* | *MC1R* regulates tyrosinase levels affecting coat colour. | [4] |
| Blanco Orejinegro, Hartón del Valle | Heat tolerance | *PRLR* | Mutations in *PRLR* cause the "slick" phenotype, causing increased thermotolerance and milk yields under tropical conditions. | [5-12] |
| All breeds | Heat tolerance | *MVD* | Identified to be under selection in North African cattle; may confer thermotolerance. | [13] |
| Limonero | Heat tolerance | *CCM2* | Involved in blood vessel development and morphogenesis; might aid heat stress adaptation. | [13] |
| Blanco Orejinegro, Hartón del Valle | Heat tolerance | *SENS2* | Downregulated in heat-stress response, helps combat protein denaturation from heat-induced stress. | [14, 15] |
| Limonero | Heat tolerance | *FBXO4* | Involved in regulation of body temperature through interactions with Hsp20. | [16, 17] |
| Blanco Orejinegro | Heat tolerance | *DNAJA2* | Part of the Hsp40 family, upregulated in response to heat stress. | [18] |
| All breeds | Heat tolerance | *HIGD1A, CBFA2T3* | Associated with cellular responses to hypoxia, could indicate a response to oxidative stress in heat-stressed cows. | [19, 20] |
| Blanco Orejinegro | Fertility and reproduction | *CATSPER3* | Exclusively expressed in the testis and encodes a sperm-specific ion channel, linked to sperm function and male fertility. | [21] |
| Blanco Orejinegro | Fertility and reproduction | *SPEF2* | Involved in the formation and functionality of sperm flagella. | [22] |
| Blanco Orejinegro, Hartón del Valle | Fertility and reproduction | *CREM, MISP3* | Implicated in spermatogenesis. | [23, 24] |
| Hartón del Valle | Fertility and reproduction | *HKDC1* | Shows high expression in the testes. | [25] |
| Limonero | Fertility and reproduction | *FSIP1* | Crucial for normal spermiogenesis and flagella development. | [26] |
| Limonero | Fertility and reproduction | *PTGES* | Encodes an enzyme involved in the synthesis of prostaglandin E2 (PGE2), a signalling molecule playing a crucial role in reproductive processes. | [27] |
| All breeds | Immune response | *CBFA2T3* | Plays a role in the cattle immune response to mammary gland inflammation. | [28] |
| Blanco Orejinegro, Limonero | Immune response | *CXCL14* | Its regulation by mRNAs and miRNAs plays a role in the immune response to bovine mastitis. | [29] |
| Limonero | Immune response | *LRRC8A* | Essential for both the development and function of T lymphocytes. | [30] |
| Blanco Orejinegro, Hartón del Valle | Resistance to anaemia | *EPB41* | Associated with resistance to anaemia, encodes proteins that form the skeletal structure of red blood cells and is linked to haematological disorders in humans. | [31] |
| Blanco Orejinegro, Limonero | Resistance to anaemia | *RPS20, RPS27* | Mutations in both genes have been shown to be responsible for Diamond-Blackfan anaemia. | [32, 33] |
| Blanco Orejinegro | Resistance to anaemia | *ZFPM1* | Plays a significant role in both cardiac and hematopoietic development, implicated in the regulation of haematopoiesis in *Trypanosoma congolense*-infected mice. | [34] |

**Table S5:** Composite selection signatures (CSS) cluster gene ontology (GO) term overrepresentation analysis for the Blanco Orejinegro breed.

| GO source | Term name | GO ID | FDR-$P_{adj.}$ | Term size | Query size | Intersect. size | Effect. domain size |
|---|---|---|---|---|---|---|---|
| GO:MF | immunoglobulin receptor binding | GO:0034987 | 3.06E+07 | 26 | 296 | 10 | 18,136 |
| GO:MF | antigen binding | GO:0003823 | 0.00001 | 57 | 296 | 10 | 18,136 |
| GO:MF | RAGE receptor binding | GO:0050786 | 0.00021 | 6 | 296 | 4 | 18,136 |
| GO:BP | phagocytosis, recognition | GO:0006910 | 6.33E+08 | 37 | 304 | 10 | 19,448 |
| GO:BP | B cell receptor signaling pathway | GO:0050853 | 0.00000 | 63 | 304 | 11 | 19,448 |
| GO:BP | complement activation, classical pathway | GO:0006958 | 0.00000 | 48 | 304 | 10 | 19,448 |
| GO:BP | humoral immune response mediated by circulating immunoglobulin | GO:0002455 | 0.00002 | 60 | 304 | 10 | 19,448 |
| GO:BP | phagocytosis, engulfment | GO:0006911 | 0.00002 | 59 | 304 | 10 | 19,448 |
| GO:BP | plasma membrane invagination | GO:0099024 | 0.00005 | 67 | 304 | 10 | 19,448 |
| GO:BP | positive regulation of B cell activation | GO:0050871 | 0.00006 | 87 | 304 | 11 | 19,448 |
| GO:BP | membrane invagination | GO:0010324 | 0.00008 | 73 | 304 | 10 | 19,448 |
| GO:BP | complement activation | GO:0006956 | 0.00008 | 73 | 304 | 10 | 19,448 |
| GO:BP | humoral immune response | GO:0006959 | 0.00009 | 187 | 304 | 15 | 19,448 |
| GO:BP | cell recognition | GO:0008037 | 0.00011 | 118 | 304 | 12 | 19,448 |
| GO:BP | regulation of B cell activation | GO:0050864 | 0.00015 | 123 | 304 | 12 | 19,448 |
| GO:BP | positive regulation of cellular process | GO:0048522 | 0.00068 | 4,242 | 304 | 101 | 19,448 |
| GO:BP | phagocytosis | GO:0006909 | 0.00099 | 175 | 304 | 13 | 19,448 |
| GO:BP | positive regulation of biological process | GO:0048518 | 0.00117 | 4,871 | 304 | 111 | 19,448 |
| GO:BP | immune response-activating cell surface receptor signaling pathway | GO:0002429 | 0.00117 | 208 | 304 | 14 | 19,448 |
| GO:BP | immune response-regulating cell surface receptor signaling pathway | GO:0002768 | 0.00243 | 223 | 304 | 14 | 19,448 |
| GO:BP | immunoglobulin mediated immune response | GO:0016064 | 0.00269 | 117 | 304 | 10 | 19,448 |
| GO:BP | B cell mediated immunity | GO:0019724 | 0.00269 | 117 | 304 | 10 | 19,448 |

| GO source | Term name | GO ID | FDR-$P_{adj.}$ | Term size | Query size | Intersect. size | Effect. domain size |
|---|---|---|---|---|---|---|---|
| GO:BP | defense response to bacterium | GO:0042742 | 0.00269 | 258 | 304 | 15 | 19,448 |
| GO:BP | antigen receptor-mediated signaling pathway | GO:0050851 | 0.00320 | 146 | 304 | 11 | 19,448 |
| GO:BP | response to bacterium | GO:0009617 | 0.00470 | 487 | 304 | 21 | 19,448 |
| GO:BP | B cell activation | GO:0042113 | 0.01092 | 230 | 304 | 13 | 19,448 |
| GO:BP | positive regulation of lymphocyte activation | GO:0051251 | 0.01142 | 232 | 304 | 13 | 19,448 |
| GO:BP | vesicle-mediated transport | GO:0016192 | 0.01240 | 1,175 | 304 | 36 | 19,448 |
| GO:BP | endocytosis | GO:0006897 | 0.01811 | 502 | 304 | 20 | 19,448 |
| GO:BP | immune response-activating signaling pathway | GO:0002757 | 0.02362 | 287 | 304 | 14 | 19,448 |
| GO:BP | neutrophil aggregation | GO:0070488 | 0.02515 | 2 | 304 | 2 | 19,448 |
| GO:BP | learned vocalization behavior | GO:0098583 | 0.02515 | 2 | 304 | 2 | 19,448 |
| GO:BP | mastication | GO:0071626 | 0.02515 | 2 | 304 | 2 | 19,448 |
| GO:BP | regulation of lymphocyte activation | GO:0051249 | 0.02515 | 363 | 304 | 16 | 19,448 |
| GO:BP | thorax and anterior abdomen determination | GO:0007356 | 0.02515 | 2 | 304 | 2 | 19,448 |
| GO:BP | negative regulation of saliva secretion | GO:1905747 | 0.02515 | 2 | 304 | 2 | 19,448 |
| GO:BP | hard palate morphogenesis | GO:1905748 | 0.02515 | 2 | 304 | 2 | 19,448 |
| GO:BP | adaptive immune response based on somatic recombination of immune receptors built from immunoglobulin superfamily domains | GO:0002460 | 0.02522 | 227 | 304 | 12 | 19,448 |
| GO:BP | positive regulation of leukocyte activation | GO:0002696 | 0.02637 | 263 | 304 | 13 | 19,448 |
| GO:BP | immune response-regulating signaling pathway | GO:0002764 | 0.03001 | 303 | 304 | 14 | 19,448 |
| GO:BP | positive regulation of cell activation | GO:0050867 | 0.03450 | 272 | 304 | 13 | 19,448 |
| GO:BP | membrane organization | GO:0061024 | 0.04225 | 554 | 304 | 20 | 19,448 |
| GO:BP | positive regulation of nitrogen compound metabolic process | GO:0051173 | 0.04225 | 2,405 | 304 | 58 | 19,448 |
| GO:CC | immunoglobulin complex, circulating | GO:0042571 | 2.87E+05 | 22 | 322 | 10 | 19,665 |

| GO source | Term name | GO ID | FDR-$P_{adj.}$ | Term size | Query size | Intersect. size | Effect. domain size |
|---|---|---|---|---|---|---|---|
| GO:CC | immunoglobulin complex | GO:0019814 | 1.11E+07 | 26 | 322 | 10 | 19,665 |
| GO:CC | organelle | GO:0043226 | 0.00345 | 13,089 | 322 | 248 | 19,665 |
| GO:CC | intracellular organelle | GO:0043229 | 0.01247 | 12,824 | 322 | 241 | 19,665 |
| GO:CC | side of membrane | GO:0098552 | 0.02013 | 435 | 322 | 18 | 19,665 |
| GO:CC | external side of plasma membrane | GO:0009897 | 0.02013 | 322 | 322 | 15 | 19,665 |
| GO:CC | intracellular anatomical structure | GO:0005622 | 0.02013 | 14,419 | 322 | 263 | 19,665 |

**Table S6:** Composite selection signatures (CSS) cluster gene ontology (GO) term overrepresentation analysis for the Hartón del Valle breed.

| GO source | Term name | GO ID | FDR-$P_{adj.}$ | Term size | Query size | Intersect. size | Effect. domain size |
|---|---|---|---|---|---|---|---|
| GO:MF | structural constituent of skin epidermis | GO:0030280 | 6.41E-08 | 21 | 276 | 16 | 18,136 |
| GO:BP | keratinization | GO:0031424 | 2.18E-01 | 34 | 289 | 16 | 19,448 |
| GO:BP | intermediate filament cytoskeleton organization | GO:0045104 | 3.39E-02 | 72 | 289 | 20 | 19,448 |
| GO:BP | intermediate filament-based process | GO:0045103 | 3.39E-02 | 73 | 289 | 20 | 19,448 |
| GO:BP | intermediate filament organization | GO:0045109 | 9.82E+00 | 50 | 289 | 16 | 19,448 |
| GO:BP | keratinocyte differentiation | GO:0030216 | 2.10E+06 | 106 | 289 | 17 | 19,448 |
| GO:BP | epidermal cell differentiation | GO:0009913 | 1.89E+08 | 161 | 289 | 18 | 19,448 |
| GO:BP | epidermis development | GO:0008544 | 0.00000 | 256 | 289 | 19 | 19,448 |
| GO:BP | cytoskeleton organization | GO:0007010 | 0.00001 | 1,205 | 289 | 44 | 19,448 |
| GO:BP | skin development | GO:0043588 | 0.00002 | 223 | 289 | 17 | 19,448 |
| GO:BP | epithelial cell differentiation | GO:0030855 | 0.00004 | 498 | 289 | 25 | 19,448 |
| GO:BP | epithelium development | GO:0060429 | 0.00006 | 899 | 289 | 35 | 19,448 |
| GO:BP | supramolecular fiber organization | GO:0097435 | 0.00023 | 628 | 289 | 27 | 19,448 |
| GO:BP | tissue development | GO:0009888 | 0.00107 | 1,444 | 289 | 44 | 19,448 |
| GO:BP | ureteric bud formation | GO:0060676 | 0.02615 | 7 | 289 | 3 | 19,448 |
| GO:BP | embryonic epithelial tube formation | GO:0001838 | 0.03174 | 102 | 289 | 8 | 19,448 |
| GO:BP | sno(s)RNA catabolic process | GO:0016077 | 0.03429 | 2 | 289 | 2 | 19,448 |
| GO:BP | epithelial tube formation | GO:0072175 | 0.03429 | 109 | 289 | 8 | 19,448 |
| GO:BP | organelle organization | GO:0006996 | 0.03429 | 2,783 | 289 | 64 | 19,448 |
| GO:BP | ureteric bud elongation | GO:0060677 | 0.03429 | 8 | 289 | 3 | 19,448 |
| GO:BP | olfactory placode development | GO:0071698 | 0.03429 | 2 | 289 | 2 | 19,448 |
| GO:BP | olfactory placode morphogenesis | GO:0071699 | 0.03429 | 2 | 289 | 2 | 19,448 |

| GO source | Term name | GO ID | FDR-$P_{adj.}$ | Term size | Query size | Intersect. size | Effect. domain size |
|---|---|---|---|---|---|---|---|
| GO:BP | olfactory placode formation | GO:0030910 | 0.03429 | 2 | 289 | 2 | 19,448 |
| GO:BP | anatomical structure development | GO:0048856 | 0.03539 | 4,196 | 289 | 88 | 19,448 |
| GO:CC | keratin filament | GO:0045095 | 3.64E-15 | 90 | 304 | 31 | 19,665 |
| GO:CC | intermediate filament | GO:0005882 | 1.28E-08 | 142 | 304 | 31 | 19,665 |
| GO:CC | intermediate filament cytoskeleton | GO:0045111 | 1.56E-08 | 172 | 304 | 33 | 19,665 |
| GO:CC | polymeric cytoskeletal fiber | GO:0099513 | 6.63E+04 | 524 | 304 | 36 | 19,665 |
| GO:CC | supramolecular fiber | GO:0099512 | 4.53E+05 | 675 | 304 | 38 | 19,665 |
| GO:CC | supramolecular complex | GO:0099080 | 4.53E+05 | 956 | 304 | 46 | 19,665 |
| GO:CC | supramolecular polymer | GO:0099081 | 4.78E+05 | 683 | 304 | 38 | 19,665 |
| GO:CC | cytoskeleton | GO:0005856 | 0.00010 | 1,766 | 304 | 53 | 19,665 |
| GO:CC | non-membrane-bounded organelle | GO:0043228 | 0.00031 | 4,965 | 304 | 111 | 19,665 |
| GO:CC | intracellular non-membrane-bounded organelle | GO:0043232 | 0.00031 | 4,964 | 304 | 111 | 19,665 |
| GO:CC | cornified envelope | GO:0001533 | 0.00275 | 29 | 304 | 5 | 19,665 |
| GO:CC | immunoglobulin complex, circulating | GO:0042571 | 0.01107 | 22 | 304 | 4 | 19,665 |
| GO:CC | intracellular organelle | GO:0043229 | 0.01776 | 12,824 | 304 | 225 | 19,665 |
| GO:CC | immunoglobulin complex | GO:0019814 | 0.01847 | 26 | 304 | 4 | 19,665 |
| GO:CC | organelle | GO:0043226 | 0.02203 | 13,089 | 304 | 228 | 19,665 |
| GO:CC | intracellular anatomical structure | GO:0005622 | 0.04658 | 14,419 | 304 | 245 | 19,665 |

**Table S7:** Composite selection signatures (CSS) cluster gene ontology (GO) term overrepresentation analysis for the Limonero breed.

| GO source | Term name | GO ID | FDR-$P_{adj.}$ | Term size | Query size | Intersect. size | Effect. domain size |
|---|---|---|---|---|---|---|---|
| GO:MF | immunoglobulin receptor binding | GO:0034987 | 5.54E+06 | 26 | 311 | 10 | 18,136 |
| GO:MF | antigen binding | GO:0003823 | 0.00001 | 57 | 311 | 10 | 18,136 |
| GO:MF | RAGE receptor binding | GO:0050786 | 0.00029 | 6 | 311 | 4 | 18,136 |
| GO:MF | L-glutamine aminotransferase activity | GO:0070548 | 0.00276 | 4 | 311 | 3 | 18,136 |
| GO:MF | oxidoreductase activity, acting on the aldehyde or oxo group of donors, oxygen as acceptor | GO:0016623 | 0.00276 | 4 | 311 | 3 | 18,136 |
| GO:MF | calcium-dependent protein binding | GO:0048306 | 0.01677 | 47 | 311 | 6 | 18,136 |
| GO:MF | signaling receptor binding | GO:0005102 | 0.02561 | 1,134 | 311 | 36 | 18,136 |
| GO:MF | cysteine-S-conjugate beta-lyase activity | GO:0047804 | 0.02561 | 2 | 311 | 2 | 18,136 |
| GO:BP | plasma membrane invagination | GO:0099024 | 8.58E+05 | 67 | 325 | 15 | 19,448 |
| GO:BP | phagocytosis, engulfment | GO:0006911 | 8.58E+05 | 59 | 325 | 14 | 19,448 |
| GO:BP | complement activation, classical pathway | GO:0006958 | 8.58E+05 | 48 | 325 | 13 | 19,448 |
| GO:BP | membrane invagination | GO:0010324 | 8.80E+05 | 73 | 325 | 15 | 19,448 |
| GO:BP | humoral immune response mediated by circulating immunoglobulin | GO:0002455 | 1.15E+08 | 60 | 325 | 13 | 19,448 |
| GO:BP | complement activation | GO:0006956 | 1.32E+09 | 73 | 325 | 13 | 19,448 |
| GO:BP | phagocytosis, recognition | GO:0006910 | 1.70E+09 | 37 | 325 | 10 | 19,448 |
| GO:BP | humoral immune response | GO:0006959 | 1.70E+09 | 187 | 325 | 19 | 19,448 |
| GO:BP | phagocytosis | GO:0006909 | 0.00000 | 175 | 325 | 17 | 19,448 |
| GO:BP | B cell receptor signaling pathway | GO:0050853 | 0.00000 | 63 | 325 | 11 | 19,448 |
| GO:BP | positive regulation of B cell activation | GO:0050871 | 0.00001 | 87 | 325 | 12 | 19,448 |
| GO:BP | immunoglobulin mediated immune response | GO:0016064 | 0.00002 | 117 | 325 | 13 | 19,448 |
| GO:BP | B cell mediated immunity | GO:0019724 | 0.00002 | 117 | 325 | 13 | 19,448 |

| GO source | Term name | GO ID | FDR-$P_{adj.}$ | Term size | Query size | Intersect. size | Effect. domain size |
|---|---|---|---|---|---|---|---|
| GO:BP | regulation of B cell activation | GO:0050864 | 0.00004 | 123 | 325 | 13 | 19,448 |
| GO:BP | lipoprotein metabolic process | GO:0042157 | 0.00006 | 129 | 325 | 13 | 19,448 |
| GO:BP | immune response-activating cell surface receptor signaling pathway | GO:0002429 | 0.00010 | 208 | 325 | 16 | 19,448 |
| GO:BP | immune response-regulating cell surface receptor signaling pathway | GO:0002768 | 0.00025 | 223 | 325 | 16 | 19,448 |
| GO:BP | lipid transport | GO:0006869 | 0.00031 | 314 | 325 | 19 | 19,448 |
| GO:BP | lipid localization | GO:0010876 | 0.00045 | 354 | 325 | 20 | 19,448 |
| GO:BP | cell recognition | GO:0008037 | 0.00085 | 118 | 325 | 11 | 19,448 |
| GO:BP | lymphocyte mediated immunity | GO:0002449 | 0.00101 | 224 | 325 | 15 | 19,448 |
| GO:BP | adaptive immune response based on somatic recombination of immune receptors built from immunoglobulin superfamily domains | GO:0002460 | 0.00113 | 227 | 325 | 15 | 19,448 |
| GO:BP | activation of immune response | GO:0002253 | 0.00144 | 355 | 325 | 19 | 19,448 |
| GO:BP | establishment of meiotic spindle localization | GO:0051295 | 0.00266 | 4 | 325 | 3 | 19,448 |
| GO:BP | positive regulation of biological process | GO:0048518 | 0.00266 | 4,871 | 325 | 115 | 19,448 |
| GO:BP | leukocyte mediated immunity | GO:0002443 | 0.00270 | 278 | 325 | 16 | 19,448 |
| GO:BP | immune response-activating signaling pathway | GO:0002757 | 0.00384 | 287 | 325 | 16 | 19,448 |
| GO:BP | defense response to bacterium | GO:0042742 | 0.00405 | 258 | 325 | 15 | 19,448 |
| GO:BP | B cell activation | GO:0042113 | 0.00426 | 230 | 325 | 14 | 19,448 |
| GO:BP | antigen receptor-mediated signaling pathway | GO:0050851 | 0.00426 | 146 | 325 | 11 | 19,448 |
| GO:BP | establishment of localization | GO:0051234 | 0.00603 | 3,760 | 325 | 92 | 19,448 |
| GO:BP | immune response-regulating signaling pathway | GO:0002764 | 0.00620 | 303 | 325 | 16 | 19,448 |
| GO:BP | membrane organization | GO:0061024 | 0.00691 | 554 | 325 | 23 | 19,448 |
| GO:BP | positive regulation of DNA replication | GO:0045740 | 0.00802 | 27 | 325 | 5 | 19,448 |
| GO:BP | synapse pruning | GO:0098883 | 0.00920 | 6 | 325 | 3 | 19,448 |

| GO source | Term name | GO ID | FDR-$P_{adj.}$ | Term size | Query size | Intersect. size | Effect. domain size |
|---|---|---|---|---|---|---|---|
| GO:BP | positive regulation of cellular process | GO:0048522 | 0.00953 | 4,242 | 325 | 100 | 19,448 |
| GO:BP | localization | GO:0051179 | 0.00989 | 4,302 | 325 | 101 | 19,448 |
| GO:BP | transport | GO:0006810 | 0.00996 | 3,622 | 325 | 88 | 19,448 |
| GO:BP | positive regulation of response to stimulus | GO:0048584 | 0.01449 | 1,627 | 325 | 47 | 19,448 |
| GO:BP | adaptive immune response | GO:0002250 | 0.01780 | 304 | 325 | 15 | 19,448 |
| GO:BP | positive regulation of cell activation | GO:0050867 | 0.01832 | 272 | 325 | 14 | 19,448 |
| GO:BP | response to bacterium | GO:0009617 | 0.01880 | 487 | 325 | 20 | 19,448 |
| GO:BP | neutrophil aggregation | GO:0070488 | 0.02097 | 2 | 325 | 2 | 19,448 |
| GO:BP | learned vocalization behavior | GO:0098583 | 0.02097 | 2 | 325 | 2 | 19,448 |
| GO:BP | mastication | GO:0071626 | 0.02097 | 2 | 325 | 2 | 19,448 |
| GO:BP | hard palate morphogenesis | GO:1905748 | 0.02097 | 2 | 325 | 2 | 19,448 |
| GO:BP | thorax and anterior abdomen determination | GO:0007356 | 0.02097 | 2 | 325 | 2 | 19,448 |
| GO:BP | negative regulation of saliva secretion | GO:1905747 | 0.02097 | 2 | 325 | 2 | 19,448 |
| GO:BP | endocytosis | GO:0006897 | 0.02387 | 502 | 325 | 20 | 19,448 |
| GO:BP | vesicle-mediated transport | GO:0016192 | 0.02402 | 1,175 | 325 | 36 | 19,448 |
| GO:BP | cell migration | GO:0016477 | 0.02741 | 1,141 | 325 | 35 | 19,448 |
| GO:BP | defense response | GO:0006952 | 0.02741 | 1,185 | 325 | 36 | 19,448 |
| GO:BP | organonitrogen compound metabolic process | GO:1901564 | 0.03104 | 5,570 | 325 | 121 | 19,448 |
| GO:BP | positive regulation of leukocyte activation | GO:0002696 | 0.03418 | 263 | 325 | 13 | 19,448 |
| GO:BP | positive regulation of lymphocyte activation | GO:0051251 | 0.03666 | 232 | 325 | 12 | 19,448 |
| GO:BP | defense response to other organism | GO:0098542 | 0.04058 | 818 | 325 | 27 | 19,448 |
| GO:BP | positive regulation of immune response | GO:0050778 | 0.04740 | 497 | 325 | 19 | 19,448 |
| GO:BP | protein metabolic process | GO:0019538 | 0.04898 | 4,814 | 325 | 106 | 19,448 |

| GO source | Term name | GO ID | FDR-$P_{adj.}$ | Term size | Query size | Intersect. size | Effect. domain size |
|---|---|---|---|---|---|---|---|
| GO:BP | vestibulocochlear nerve formation | GO:0021650 | 0.04978 | 3 | 325 | 2 | 19,448 |
| GO:BP | zygotic determination of anterior/posterior axis, embryo | GO:0007354 | 0.04978 | 3 | 325 | 2 | 19,448 |
| GO:CC | immunoglobulin complex, circulating | GO:0042571 | 3.85E+05 | 22 | 332 | 10 | 19,665 |
| GO:CC | immunoglobulin complex | GO:0019814 | 1.49E+07 | 26 | 332 | 10 | 19,665 |
| GO:CC | extracellular region | GO:0005576 | 0.01611 | 1,632 | 332 | 48 | 19,665 |
| GO:CC | complement component C1q complex | GO:0062167 | 0.03083 | 2 | 332 | 2 | 19,665 |

**Table S8:** Composite selection signatures (CSS) cluster QTLs identified for all breeds [Excel file – Ward_et_al.(2023)_Table_S8.xlsx].

### SUPPLEMENTARY MATERIAL REFERENCES
